# Profiling epigenetic age in single cells

**DOI:** 10.1101/2021.03.13.435247

**Authors:** Alexandre Trapp, Csaba Kerepesi, Vadim N. Gladyshev

**Affiliations:** Division of Genetics, Department of Medicine, Brigham and Women’s Hospital, Harvard Medical School, Boston, MA 02115, USA

## Abstract

DNA methylation of a defined set of CpG dinucleotides emerged as a critical and precise biomarker of the aging process. Multi-variate machine learning models, known as epigenetic clocks, can exploit quantitative changes in the methylome to predict the age of bulk tissue with remarkable accuracy. However, intrinsic sparsity and digitized methylation in individual cells have so far precluded the assessment of aging in single cell data. Here, we present scAge, a probabilistic approach to determine the epigenetic age of single cells, and validate our results in mice. scAge tissue-specific and multi-cell type single cell clocks correctly recapitulate chronological age of the original tissue, while uncovering the inherent heterogeneity that exists at the single-cell level. The data suggest that while tissues age in a coordinated fashion, some cells age more or less rapidly than others. We show that individual embryonic stem cells exhibit an age close to zero, that certain stem cells in a tissue show a reduced age compared to their chronological age, and that early embryogenesis is associated with the reduction of epigenetic age of individual cells, the latter supporting a natural rejuvenation event during gastrulation. scAge is both robust against the low coverage that is characteristic of single cell sequencing techniques and is flexible for studying any cell type and vertebrate organism of interest. This study demonstrates for the first time the potential for accurate epigenetic age profiling at single-cell resolution.

## INTRODUCTION

Among several canonical hallmarks, aging involves profound epigenetic alterations, particularly at CG dinucleotides (CpGs) (Horvath & Raj, 2018; López-Otín et al., 2013). Changes in CpG methylation with age can now be assayed using a variety of approaches, ranging from hybridization arrays to genome-wide or targeted next-generation sequencing methods (Han, Franzen, et al., 2020; Horvath, 2013; Lister et al., 2009; Meissner et al., 2005). These techniques permit quantitative examination at single-base resolution of the dynamic DNA methylation landscape in any tissue of interest in organisms, such as mammals, that evolved this type of regulation.

Since their inception in the last decade, predictive multi-variate machine learning models based on DNA methylation (DNAm) levels, termed ‘epigenetic clocks,’ have revolutionized the aging field (Bocklandt et al., 2011; Horvath, 2013). First built strictly as an estimator of chronological age, clocks can now also integrate and predict various measures of biological aging and disease risk, underscoring their clinical relevance (Levine et al., 2018; Lu et al., 2019). Excitingly, several pan-tissue mammalian clocks were recently developed that can profile epigenetic age in virtually any tissue across eutherians with impressive precision (Mammalian Methylation Consortium, 2021). On top of these remarkable advancements, epigenetic clocks are of particular interest within the scopes of lifespan extension and cell reprogramming, as these models show promise in detecting even small changes in biological age that result from these interventions (Lu et al., 2020; Petkovich et al., 2017).

However, while individual cells are the units of life, all existing epigenetic clocks rely on measurements derived from bulk samples (i.e., samples containing many cells), both for the creation and application of these models (Bell et al., 2019). Historically, using bulk samples for DNA methylation analysis has been an inherent requirement of the methodologies available, which demanded hundreds of nanograms of input material due to harsh chemical treatment of DNA by bisulfite conversion (Karemaker & Vermeulen, 2018). While the use of bulk tissue is convenient, it also inherently obscures the epigenetic heterogeneity that exists among individual cells (Bell et al., 2019; Gravina et al., 2016). Recent work was done to characterize the transcriptomic changes in murine aging at single-cell resolution, but detailed and cross-tissue single-cell epigenetic changes during aging in mammals remain mostly unexplored (Almanzar et al., 2020).

Recent advances in epigenomic sequencing methods have made it possible to evaluate limited methylation profiles in single cells. Since the inception of these techniques in the previous decade, a variety of single-cell methylation sequencing methods have surfaced, including single-cell reduced representation (scRRBS) and single-cell (whole genome) bisulfite sequencing (scWGBS/scBS) (Gravina et al., 2016; Guo et al., 2013; Smallwood et al., 2014). These approaches rely on key modifications of the original bulk techniques that enable reduced DNA loss during library preparation. Excitingly, novel methods for sequencing RNA, DNA methylation, and chromatin accessibility in the same single cell have also been devised, allowing for integration of multi-omic analyses at maximal resolution (Angermueller et al., 2016; Argelaguet et al., 2019; Clark et al., 2018).

Despite this remarkable progress in single-cell omics, there remain common issues of sparsity. Depending on the particular method used, only a small fraction of CpGs covered with bulk sequencing methods are represented in single cells (Fig. 1a, Fig. S1). Furthermore, the most widespread protocols for single cell methylome profiling—those involving genome-wide interrogation of DNA methylation patterns—suffer additionally from effectively random coverage of reads (Karemaker & Vermeulen, 2018). To overcome this limiting factor, analysis of single cell methylation profiles is usually conducted by averaging methylation levels in genomic bins (Angermueller et al., 2016; Luo et al., 2017). Alternatively, several imputation and clustering strategies have also been developed, employing Bayesian or deep learning approaches to fill-in missing methylation states for CpGs not covered in a given cell (Angermueller et al., 2017; Kapourani & Sanguinetti, 2019). While these imputation methods work very well to distinguish cellular subtypes from one another, they rely on building dataset-specific models, rendering it difficult to perform unbiased comparisons between studies.

**Figure 1:**
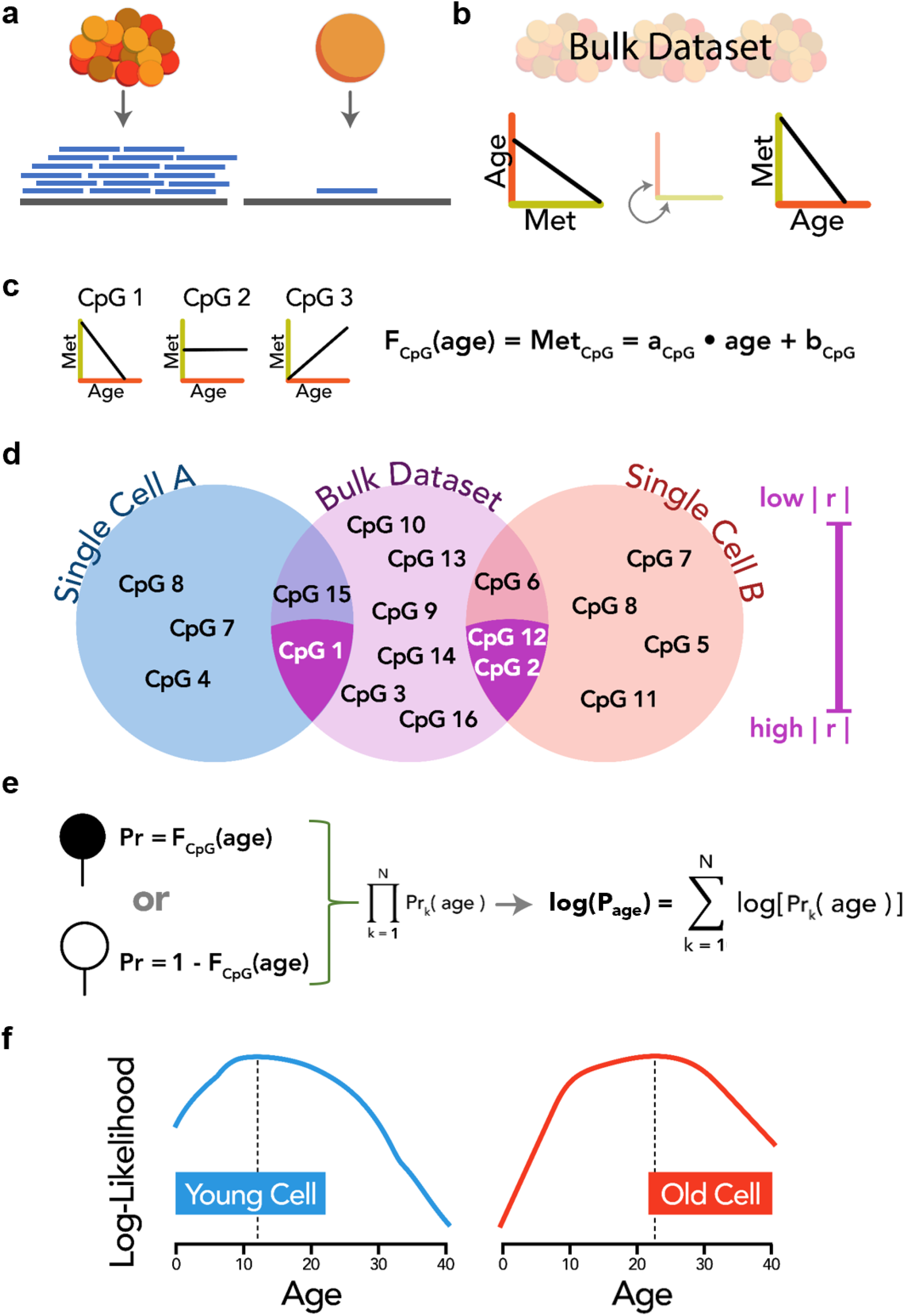
Developing scAge. **a)** Schematic of differential read alignment with bulk (left) and single-cell (right) epigenomic sequencing methods. Decreased DNA input in single cells results in fewer mapped reads (blue) onto a reference genome (grey), leading to decreased CpG coverage compared to bulk samples. **b)** Schematic reversion in the linear relationship between methylation level and age. Using bulk samples of different ages, it is possible to map how methylation level (green axis) influences or predicts age (red axis). Conversely, the opposite relationship can also be established, whereby age can be used as a predictor of average methylation level. **c)** Computation of linear regression formulas for every CpG in the training dataset. CpG methylation level can increase, decrease, or have no apparent linear relationship with age. *Met* denotes the average methylation level, a indicates the slope of the linear equation, and *b* represents the intercept along the methylation axis. **d)** Schematic intersection and filtering step of scAge. CpGs in the training dataset (pink) are intersected with those in individual cells (blue/orange). CpGs in one single cell show limited intersections with those in another single cell. Common CpGs between any one cell and the bulk training dataset are ranked based on their absolute correlation |r| with age. Only the top correlated common CpGs (dark purple intersections) in each cell are taken as input to the scAge algorithm. **e)** Schematic of the core scAge algorithm. Posterior probabilities are computed based on observed methylation states in single cells for every age step. Filled-in circle indicates a methylated CpG, while an open circle indicates an unmethylated CpG in a particular single cell. A likelihood metric for the profile is computed as the product of all independent probabilities per CpG. Practically, this is evaluated as a log-likelihood sum to overcome computational underflow limitations. **f)** Schematic representation of the distribution of age-dependent log-likelihood metrics in a young and old cell.

The overall sparsity in single-cell DNAm profiles poses profound limitations for the creation of single-cell epigenetic clocks. Building these predictive models traditionally relied on collecting methylation levels of CpGs covered consistently between samples of different ages (Meer et al., 2018; Stubbs et al., 2017; Thompson et al., 2018). In bulk tissue, this enables the creation of large feature tables that can then be directly harnessed for machine learning, particularly elastic net regression (Zou & Hastie, 2005). However, sparse and binarized methylation profiles of single cells preclude this approach (Fig. S1) (Bell et al., 2019). Despite these challenges, the creation of single-cell epigenetic clocks promises both new methods for ultra-low-input epigenetic age profiling, combined with unparalleled resolution into the aging process.

Here, we developed scAge, a novel clock method capable of profiling epigenetic age in single cells. Due to low and inconsistent CpG coverage, our approach instead relies on a probabilistic algorithm that is largely independent of which CpGs are covered in each cell. By harnessing the linear relationship of methylation levels with age in a subset of CpGs, we create a likelihood metric that quantifies the probability of a cell to come from a bulk sample of a given age. Our method recapitulates the chronological age of the tissue on average, while also uncovering the intrinsic epigenetic heterogeneity that exists between cells. Use of these probability-based epigenetic clocks opens up exciting new avenues for research on biological aging at the previously elusive level of single cells.

## RESULTS

### Designing scAge: a probability-based single cell epigenetic clock

Major challenges in assessing the age of single cells are their sparse and binarized methylation profiles. Contrary to bulk samples, the sequence reads cover different parts of the genome of each single cell with very low overlap among the cells (Fig. 1a, Fig. S1). To overcome these limitations, we assumed that the methylation levels of highly covered CpG sites in bulk sequencing or DNAm array profiling of a tissue offer an estimation of the probability of methylation at these particular CpG sites in any single cell coming from that tissue. Additionally, we reversed the conventional notion about the relationship between methylation level and age: while current bulk clocks use methylation level as a predictor of age, we hypothesized that age could be thought of as a predictor of bulk methylation level at any given CpG (Fig. 1b). Using training data derived from bulk RRBS, we estimated the change in average methylation levels with age for each CpG in this reference set using univariate linear models (Fig. 1c).

Next, we isolated the common CpG sites between any given single cell profile and the reference methylation probability dataset (Fig. 1d). We then selected a defined number of common CpGs that exhibited the greatest absolute Pearson correlation with age in the bulk data (i.e., the age-associated CpG sites). Importantly, due to the sparsity of single cell DNAm profiles, covered CpG sites vary greatly from cell to cell; despite this, a small but distinct collection of age-associated CpG sites are covered in every sequenced cell (Fig. 1d).

Then, we calculated the likelihood of observing this filtered methylation profile of an individual cell at any given age (Fig. 1e). Practically, we applied logarithms (log-likelihood) to avoid underflow errors during computation. Finally, we determined the age for which this likelihood is maximal (Fig. 1f). We found that this approach, which we designate scAge, permitted accurate epigenetic age profiling in single cells with dramatically different and sparse methylome profiles.

### Probabilistic single cell clocks recapitulate the chronological age of single hepatocytes

We first applied scAge to terminally differentiated cells from very young and very old mice, including 11 single hepatocytes from 4-month-old animals and 10 single hepatocytes from 26-month-old animals (Gravina et al., 2016). Single-cell profiles contained limited common CpGs between any given pair; in fact, this effect was greatly accentuated when sites in additional cells were progressively intersected, resulting in minimal final overlap (Fig. 2a). The coverage ranged from 0.4-3.2 million CpGs in hepatocytes, with similar mean global methylation in young and old cells (Fig. 2b, Fig. S2a). We first applied our probabilistic clock trained on bulk liver samples (Fig. 2c) (Thompson et al., 2018). Remarkably, using only 700 independent CpGs per cell, scAge showed both impressive accuracy and consistency in age predictions for young and old hepatocytes (Fig. 2d). We achieved a Pearson r coefficient of 0.88 with mean and median absolute errors of 3.9 and 2.9 months, respectively. Thus, liver scAge correctly recapitulated the age of the original tissue with only a handful of cells.

**Figure 2:**
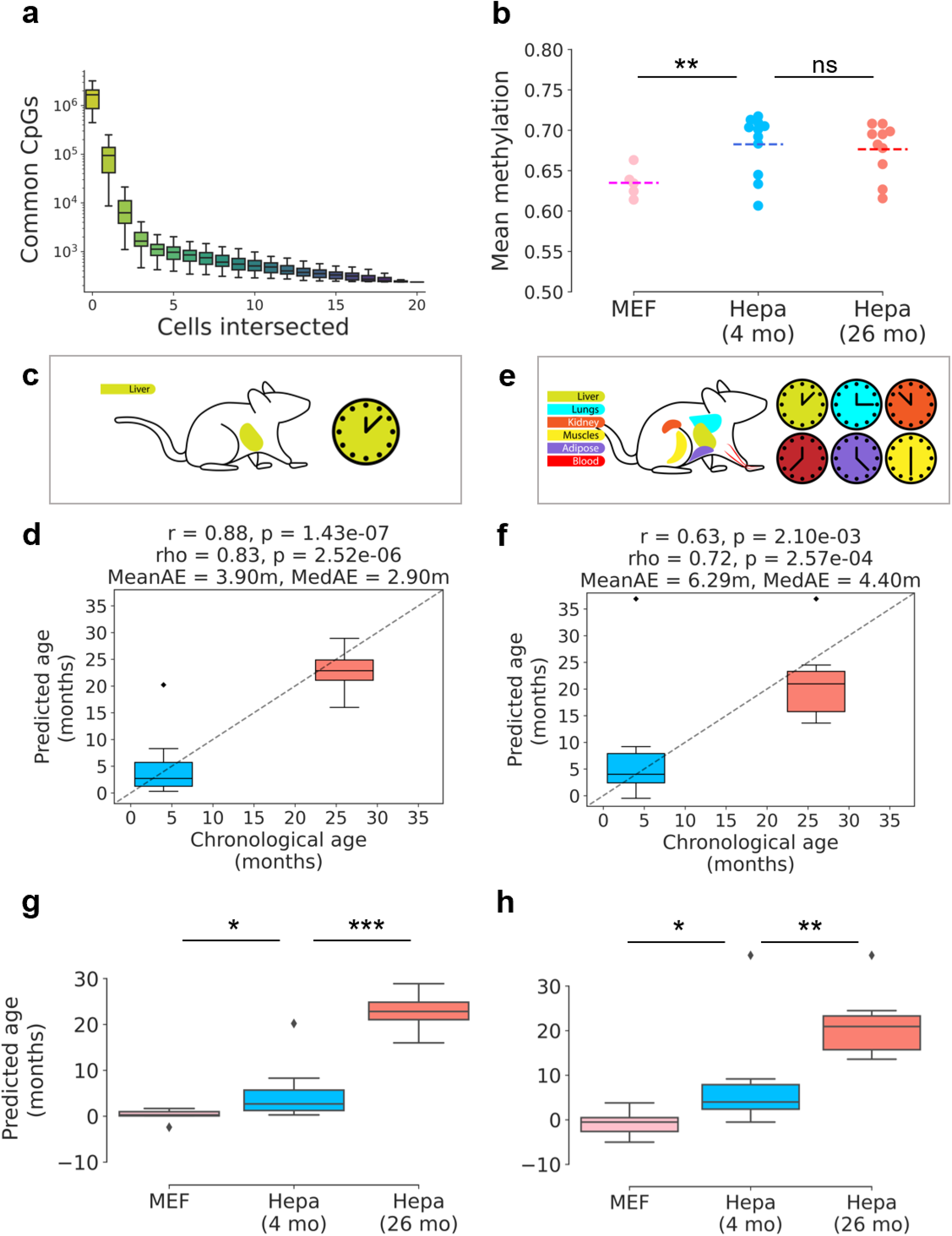
scAge recapitulates the age of hepatocytes and embryonic fibroblasts. **a)** Progressive intersection of CpG sites covered in each individual hepatocyte. The order of intersections was permuted 100 times to produce distributions for every additional cell added. Y-axis is log-transformed. Minimal common CpGs remain after all cells were intersected. **b)** Mean methylation in MEFs, 4-month-old and 26-month-old hepatocytes. ** indicates p < 0.01. **c)** Schematic representation of the liver scAge clock. **d)** Epigenetic age predictions for all young and old hepatocytes using the liver scAge clock. **e)** Schematic representation of the multi-tissue scAge clock. Tissues included were liver, lung, kidney, muscle, adipose, and blood. **f)** Epigenetic age predictions for all young and old hepatocytes using the multi-tissue scAge clock. Application of the liver scAge clock **(g)** and the multi-tissue scAge clock **(h)** on mouse embryonic fibroblasts (MEFs) compared to young and old hepatocytes. * denotes p < 0.05, ** denotes p < 0.01, and *** denotes p < 0.001.

While scAge serves well to integrate the predictions for multiple single cells into an accurate predictor of overall tissue age, it also inherently provides increased resolution down to individual cells. Indeed, some cells were predicted to be younger or older than others in the same tissue. The lowest prediction using the liver clock for young cells was close to 0, while one cell was predicted to be around 20 months old. These results suggest deep heterogeneity in the aging process, wherein global changes in epigenetic age in a bulk tissue are characterized by the uneven and diverse aging trajectories that individual cells undergo.

We further employed scAge trained on a multi-tissue dataset consisting of kidney, blood, liver, lung, muscle, and adipose tissues (Fig. 2e) (Thompson et al., 2018). Because multi-tissue datasets add biological noise to the relationship between age and methylation level at most CpGs, absolute correlations between both variables dropped drastically compared to a liver-exclusive dataset (Fig. S3). Due to this, we reasoned that predictive metrics would improve using a multi-tissue dataset if more CpGs were considered per cell to compute age likelihoods. Thus, we used the multi-tissue scAge predictor with 2,000 CpGs profiled per cell. This model showed decreased accuracy compared to the liver model, with a Pearson *r* of 0.63 (Spearman rho = 0.72) and mean and median absolute errors of 6.29 and 4.4 months, respectively (Fig. 2f). Interestingly, the multi-tissue model predicted the age of one cell in each group to be near the maximum age that we designated when running the algorithm. We interpret these observations as an accelerated aging trajectory (i.e., accelerated senescence) of some cells from the population, further underscoring the heterogeneity of epigenetic aging in single cells. Removing both outliers in the liver and multi-tissue models resulted in much improved prediction accuracy, with Pearson r values of 0.95 and 0.9, respectively (Fig. S4).

Interestingly, the predictive metrics of these two models on the liver data varied based on the number of CpGs included in the overall likelihood calculation, whereby incorporating too few or too many CpGs resulted in decreased prediction accuracy (Fig. S5). When too few CpGs were used, there were not enough individual probabilities to compute a precise age prediction (Fig. S5a-b). However, the inclusion of too many CpGs also led to a slight decrease in the predictive accuracy of the models (Fig. S5c-d). Because our algorithm ranked CpGs based on how they are correlated with age, we suggest that including more lower-ranked CpGs introduced noise into the prediction, thereby decreasing overall accuracy. Of note, the accuracy of predictions did not show a significant relationship with the number of common CpGs between any one single cell and the training dataset (Fig. S6). This implies that our method is robust to relatively low coverage in single cells.

### scAge clocks predict the age of embryonic fibroblasts close to zero

We also applied scAge to 5 mouse embryonic fibroblasts (MEFs) included in the same dataset. MEFs had significantly higher coverage than the hepatocytes, owing to the improved DNA quality that resulted from a milder isolation process compared to the liver cells (Fig. S2a) (Gravina et al., 2016). Additionally, the mean methylation in MEFs was lower than that of the hepatocytes (Fig. 2b). scAge trained on either the liver or the multi-tissue datasets predicted the epigenetic age of MEFs to be around 0 (Fig. 2g-h). Despite being in culture, these cells appeared to retain the epigenetic age information from the embryo. Overall, our results show that scAge, whether trained on a liver or multi-tissue dataset, can accurately recapitulate the chronological age of the tissue of origin in single hepatocytes and embryonic fibroblasts.

### Muscle stem cells show minimal epigenetic aging

To further investigate the applicability of scAge, we applied it to young and old muscle stem cell data (Hernando-Herraez et al., 2019). This dataset consisted of 275 single cells from 6 donors, including 4 young (1.5 months) and 2 old animals (26 months). Due to technical variability in the scBS methodology, only 185 cells (67%) had greater than 1 million CpGs covered (Fig. S2b). Mean methylation between young and old cells was comparable.

When we applied the same multi-tissue scAge that profiles 2,000 CpGs to these muscle stem cells, the epigenetic age of young cells was 9.5 weeks on average, roughly concordant with their chronological age. Interestingly, old muscle stem cells showed a significant epigenetic age increase, but only on the order of a few weeks, with a mean predicted age of 18.3 weeks (Fig. 3a-b). These results are coherent with the previous analysis that examined epigenetic age of these cells via a pseudo-bulk grouping approach utilizing a muscle clock trained with conventional elastic net methods (Hernando-Herraez et al., 2019; Reizel et al., 2015; Stubbs et al., 2017). Overall, our results agree with the previously reported epigenetic aging dynamics of mouse muscle stem cells, but offer single-cell resolution to the data.

### Single embryonic stem cells display low epigenetic age

We next sought to evaluate scAge on the most common type of publicly available single-cell methylation datasets: those profiling embryonic tissue. Embryonic stem cells (ESCs) and their induced pluripotent stem cell (iPSC) counterparts generally show very low predicted epigenetic ages trending towards 0 (Horvath, 2013; Meer et al., 2018; Petkovich et al., 2017). To test our model, we examined 3 datasets of embryonic stem cells and related tissues (Angermueller et al., 2016; Clark et al., 2018; Smallwood et al., 2014). Cells from these studies showed variable coverage, and we selectively filtered those that had at least 1 million CpGs covered to improve consistency between datasets (Fig. S2c). Importantly, ESCs were cultured in either traditional serum conditions, or grown in serum-free media supplemented with the “2i” cocktail of MEK and GSK3β inhibitors. Culturing cells in 2i medium was previously shown to drive global hypomethylation in ESCs, producing epigenetic profiles concordant with migrating primordial germ cells (Ficz et al., 2013).

As expected, we observed significant hypomethylation among 2i cells in both the Angermueller et al. and Smallwood et al. studies (Fig. 3c). MII oocytes included in the Smallwood et al. study showed comparable mean global methylation to 2i cells, and embryoid bodies derived from ESCs in the Clark et al. study had the greatest mean methylation on average, significantly higher than both serum-grown and 2i ESCs (Fig. 3c). We applied our liver and multi-tissue scAge models on all filtered cells from these three studies, and observed consistently low predicted ages for all cell types assayed (Fig. 3d-e). Using the liver model, ESCs cultured in serum displayed an epigenetic age around 0, while 2i ESCs showed significantly higher predicted epigenetic age (Fig. 3d). However, the multi-tissue scAge predictor showed a contrasting trend (Fig. 3e). Additionally, the multi-tissue model showed greater variance and more extreme predictions compared to the liver model in all three datasets. We suggest that this may occur due to multi-tissue datasets depicting less robust linear relationships with age, whereby the increased variation in methylation levels between different tissues effectively translates to less consistent predictions (Fig. S3). Embryoid bodies derived from serum-grown ESCs demonstrated higher age using both clocks, hinting that the initiation of unmodulated differentiation signals into the three germ layers rapidly induces a noticeable increase in the epigenetic age of cells. Overall, our results predicted embryonic stem cells to be close to 0 in epigenetic age, while uncovering significant differences based on particular culture conditions.

**Figure 3:**
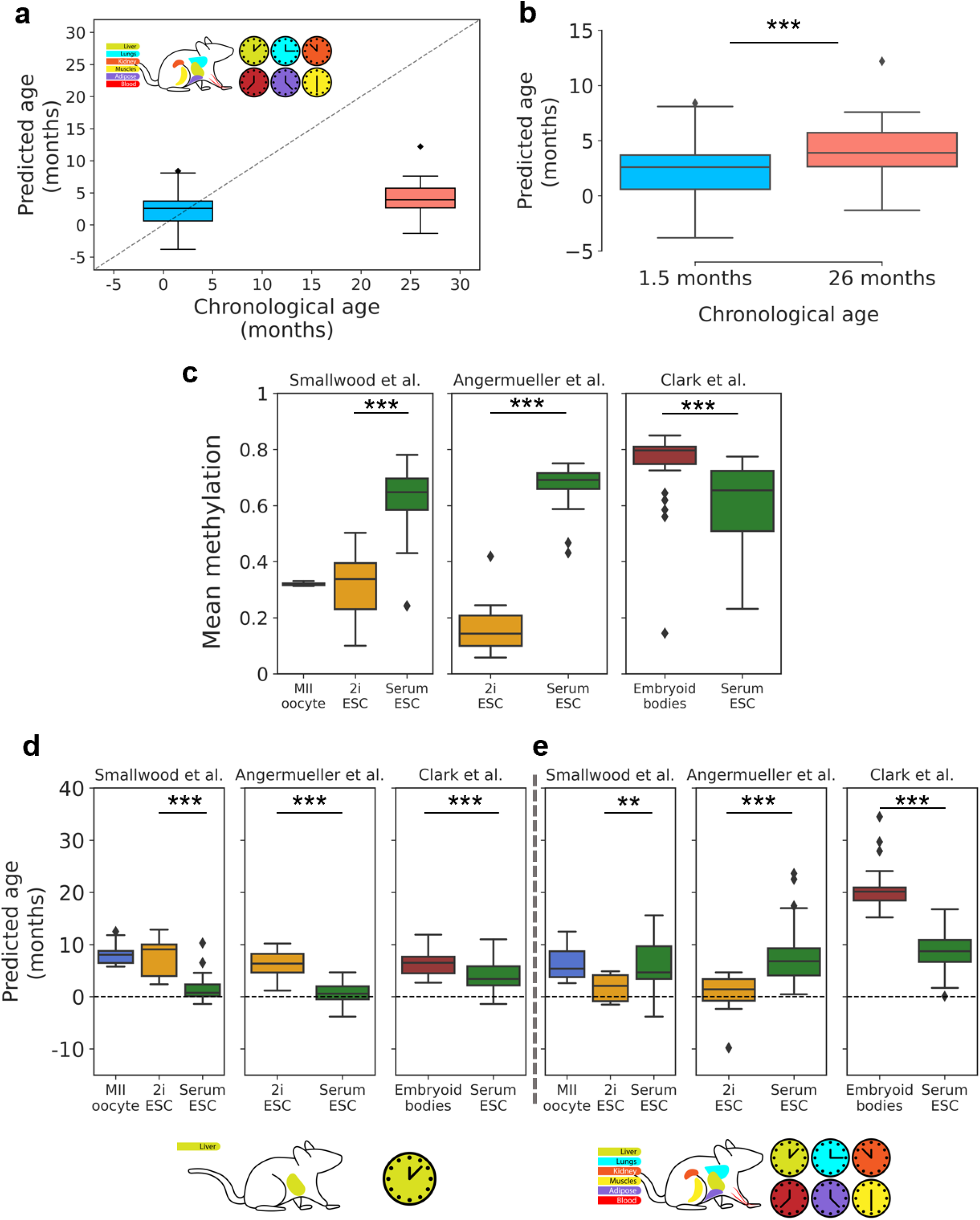
scAge reveals distinct epigenetic aging trajectories in stem cells. Epigenetic age predictions of muscle stem cells using the multi-tissue scAge clock plotted against chronological age **(a)** or side-by-side **(b)**. *** denotes p & 0.001. **c)** Mean methylation of oocytes, embryonic stem cells (ESCs) in 2i or serum culture conditions, and embryoid bodies. Cells are grouped by the study of origin. *** denotes p < 0.001. Epigenetic age predictions in all cell types assayed in the three embryonic studies by the liver scAge clock **(d)** or the multi-tissue scAge clock **(e)**. ** denotes p & 0.01, *** denotes p < 0.001.

### Single cell analyses suggest a rejuvenation event during mouse gastrulation

We then investigated a dataset profiling mouse gastrulation at single-cell resolution (Argelaguet et al., 2019). This data consisted of 758 single cells isolated from murine embryos ranging from embryonic day (E) 4.5 to 7.5. We filtered the data to keep single cells with at least 500,000 CpGs covered, resulting in a final dataset of 495 cells (Fig. S2d). Interestingly, mean global methylation varied drastically during this early period of mouse gastrulation, with E4.5 cells showing significant hypomethylation compared to the other three developmental stages (Fig. 4a). This trend in global methylation suggests a link between ESCs grown in 2i conditions and single cells from E4.5 embryos.

**Figure 4:**
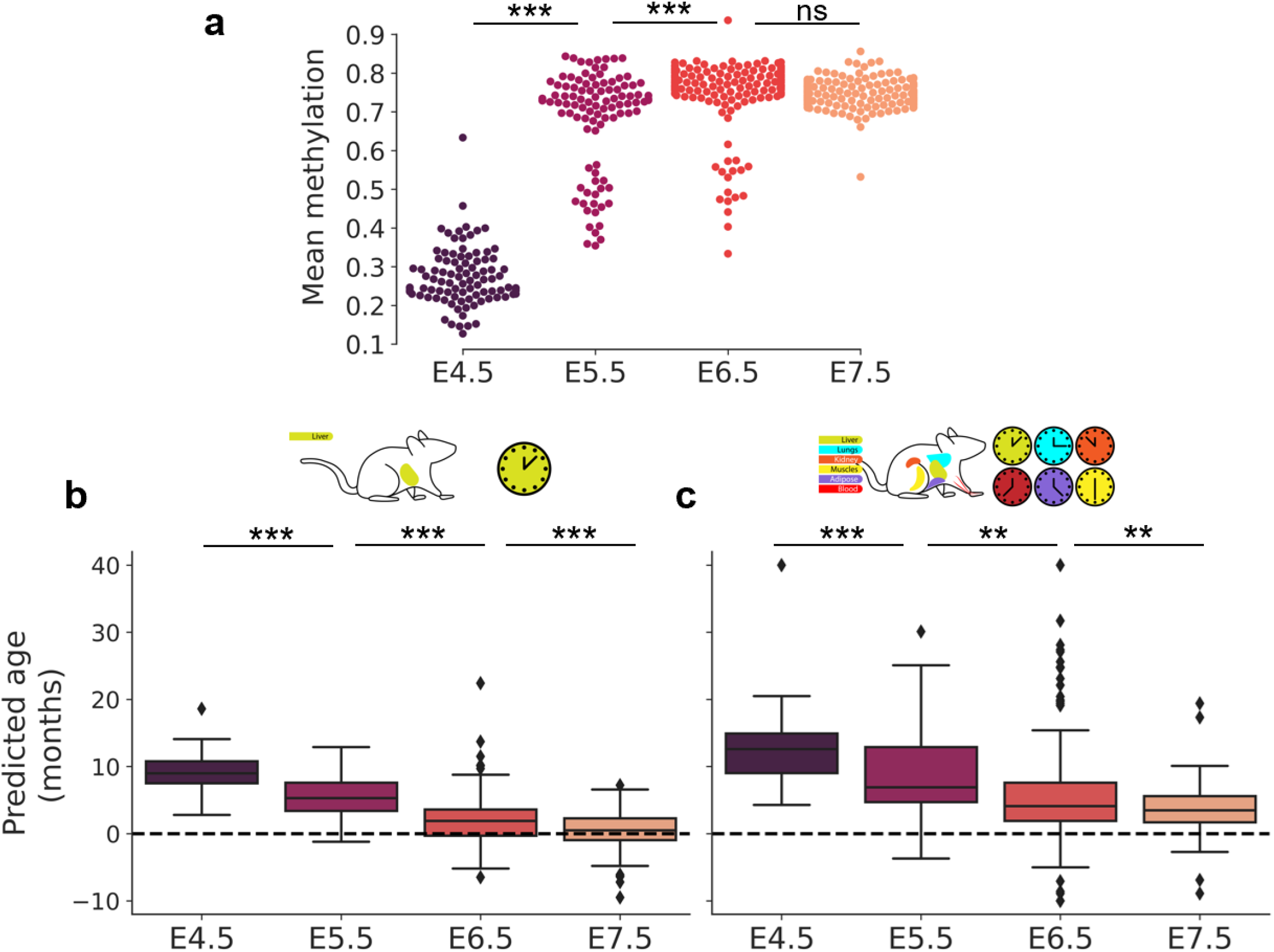
scAge suggests a rejuvenation event during gastrulation. **a)** Mean methylation among E4.5 to E7.5 cells. *** denotes p < 0.001. Epigenetic age predictions for single embryonic cells throughout gastrulation with the liver scAge clock **(b)** and the multi-tissue scAge clock **(c)**. ** denotes p < 0.01, *** denotes p < 0.001.

It was recently suggested that embryogenesis may be characterized by an initial decrease in biological age to a point termed the “ground zero,” after which organismal aging formally begins (Gladyshev, 2021). Consistent with this idea, recent application of epigenetic clocks to bulk samples revealed a significant decrease in biological age (i.e., rejuvenation) during early stages of embryogenesis, followed by an increase in later stages (Kerepesi et al., 2021). This finding also agrees with the notion that damage accumulation inevitably occurs during the lifespan of an organism, even in germ cells. Thus, a rejuvenation event is thought to take place during mid-embryogenesis to ensure the continuous generation of new biologically young individuals.

To investigate this idea at the single-cell level, we applied both the liver and multi-tissue scAge clocks to single embryonic cells from the four developmental stages assayed. Liver scAge showed a steady and significant reduction in the mean predicted age from E4.5 to E7.5, with the latter reporting the age around 0 (Fig. 4b). The multi-tissue scAge clock showed an identical trend, albeit with slightly elevated and more variable predicted ages (Fig. 4c). Together, these results show that a rejuvenation event occurs during mid-embryogenesis and that individual cells may be rejuvenated through natural means. Notably, the lowest single-cell epigenetic age approximately corresponds to the stage of gastrulation and is associated with hypermethylation, suggesting that to rejuvenate cells it is important to both demethylate and subsequently remethylate the genome.

## DISCUSSION

In this work, we report scAge, a probabilistic model to ascertain the epigenetic age of single cells. Our method utilizes bulk methylation data to train linear regression models that predict methylation levels given exclusively age as the input. Based on these univariate models, we compute the posterior probability of observing an unmethylated or methylated state in a single cell. Using a selected fraction of age-related CpGs and their associated probabilities, we calculate the likelihood that a cell comes from a tissue of a certain chronological age and register the age of maximum likelihood as our ultimate predictor of epigenetic age. This approach solves the challenges of sparsity and uneven coverage of methylation profiles of single cells, which precluded attempts to estimate epigenetic age in individual cells. Indeed, all previous epigenetic clocks require defined sets of CpG sites for their application, which is not feasible in the case of single cells.

This method enables accurate age prediction of single hepatocytes and mouse embryonic fibroblasts with high resolution on models trained either on liver or multi-tissue datasets. Additionally, we show consistency between our model and previous work in mouse muscle stem cells, which display attenuated epigenetic aging in comparison to their chronological age. We also find that while ESCs are generally predicted to have low epigenetic age, the age differs depending on the culture condition. Finally, our data provide further evidence for the “ground zero” hypothesis of aging by showing a highly significant and steady decrease in the epigenetic age of single cells at the time of gastrulation.

Despite this advance in epigenetic age profiling in single cells, various avenues for improvement exist. For one, binary methylation states of CpGs were here assumed to be independent of each other, as prior work suggested that this was the case when analyzing single reads from bulk samples (Han, Franzen, et al., 2020; Han, Nikolić, et al., 2020). However, a more thorough analysis of this behavior specifically in single cells may reveal biological insights suggesting a more complex relationship. Additionally, using only linear regression may be suboptimal when considering the potentially diverse set of mathematical associations that best model CpG methylation levels and age. Lastly, it remains to be explored how the individual aging trajectories of cells change with time, and how these are transferred during events such as cell division.

Taken together, these results suggest dramatic implications regarding epigenetic aging. We find that the aggregation of multiple single cell predictions provides an accurate average measure of the age of a particular tissue. However, this single cell approach concurrently discovers profound heterogeneity in the aging trajectories of individual cells. This suggests that all cells in a tissue do age, but that clocks likely tick independently within single cells. In turn, we suggest some cells undergo accelerated or decelerated epigenetic aging, which was previously impossible to ascertain (Fig. 5a). Additionally, this method may have profound clinical applications for human somatic, germline, and cancer cells, as it may be possible to discriminate and map “young” and “old” cells within a heterogeneous tissue via this approach (Fig. 5b). Overall, we present here the first method to determine epigenetic age in single cells, with far-reaching potential across the field of aging.

**Figure 5:**
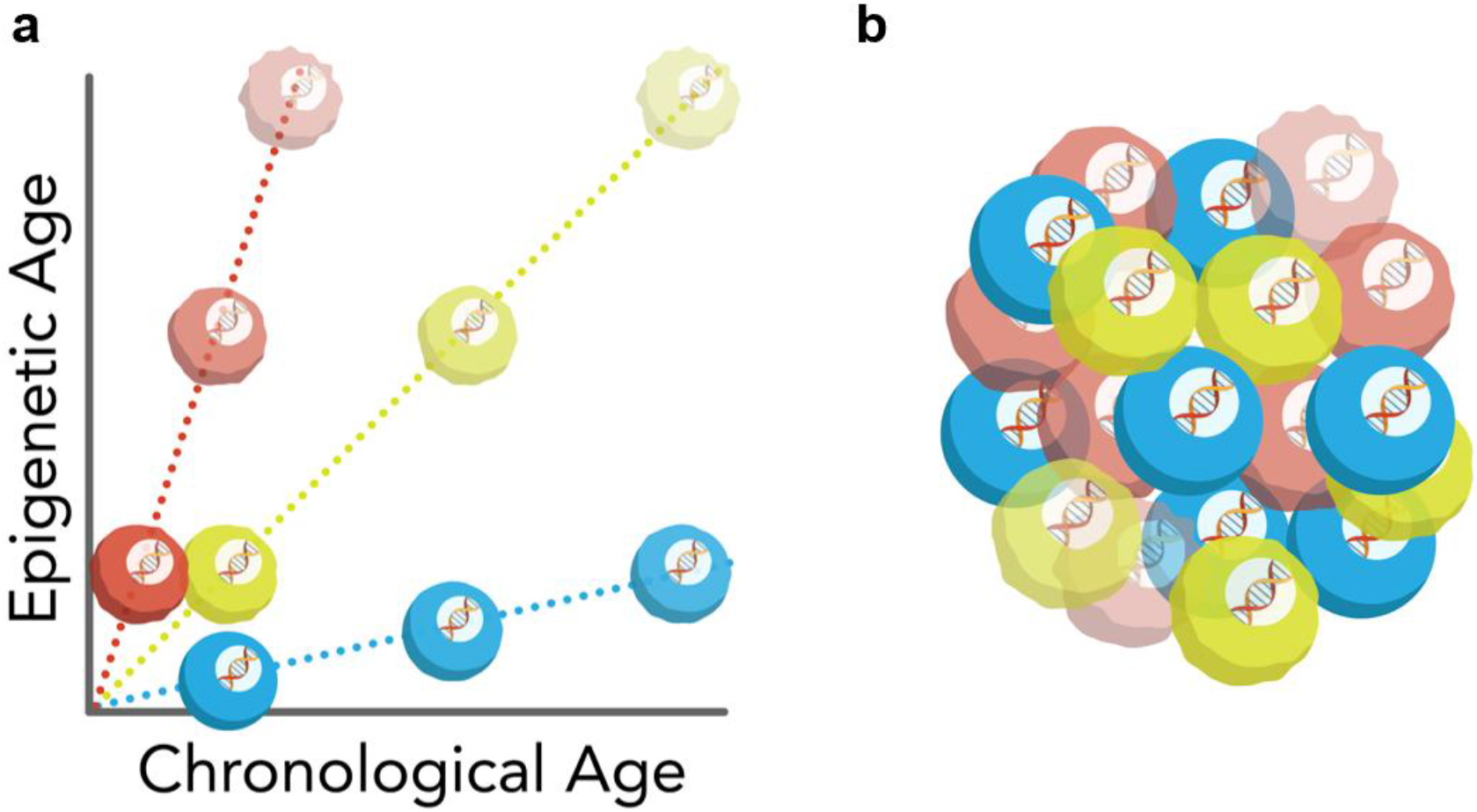
Models of aging in single cells. **a)** Schematic model of aging trajectories of single cells. Some cells (red) show accelerated epigenetic aging, while other cells (blue) show decelerated epigenetic aging compared to a reference cell population (green). **b)** Schematic representation of epigenetic aging heterogeneity within a population of cells, which is composed of cells with accelerated (red), decelerated (blue) and average (green) aging trajectories.

## METHODS

### Single cell data processing

For the Gravina et al. study, sequence data was downloaded from the SRA under accession number SRA344045 (Gravina et al., 2016). In this case, sequence data was pre-trimmed prior to deposition to the SRA. Trimmed sequences were mapped to the mm10/GRCm38.p6 genome using Bismark v0.22.3 with the option –*non_directional*, as suggested by the Bismark User Guide v0.21.0 for Zymo Pico-Methyl scWGBS library preparations. Reads were deduplicated and methylation levels for CpG sites were extracted with Bismark (Krueger & Andrews, 2011).

For the Hernando-Herraez et al., Angermueller et al., Clark et al., Smallwood et al., and Argelaguet et al. studies, processed coverage files containing extracted methylation levels generated by Bismark were downloaded directly from the GEO database under accession numbers GSE121436, GSE68642, GSE109262, GSE56879, and GSE121690, respectively (Angermueller et al., 2016; Argelaguet et al., 2019; Clark et al., 2018; Hernando-Herraez et al., 2019; Smallwood et al., 2014).

All coverage files were then further processed to scale methylation level to a ratio between [0, 1]. While single cell methylation profiles were almost entirely binary, technical considerations such as PCR amplification bias resulted in some intermediate methylation values. Uncertain methylation calls of 0.5 were removed prior to downstream analysis. Remaining methylation values were rounded to 0 or 1. Genomic positions on the 19 mouse autosomes were retained for analysis to partially minimize the effect of sex on the study. Coverage was interpreted as the total number of covered methylated and unmethylated cytosines on both DNA strands. Average methylation in single cells was computed as the mean of all binary methylation states observed.

### Bulk data processing

In order to create bulk reference datasets that estimate the linear relationship between age and methylation level, we downloaded processed RRBS data from the Thompson et al. study deposited in the GEO database under accession number GSE120132 (Thompson et al., 2018). This dataset consisted of 549 total samples from liver, lung, blood, kidney, adipose and muscle tissues with ages ranging from 1 month to 21 months. Methylation fractions were taken as the number of reads supporting a methylated status for a CpG over the total number of reads that covered this CpG. To maximize the accuracy of bulk methylation levels while also preserving as many sites as possible, only CpG sites for which 90% of samples had at least 5x coverage in were retained. This resulted in a final multi-tissue matrix of 549 samples by 748,955 positive strand CpGs (autosomic chromosomes only) with some missing values. From here, a separate liver-only matrix containing 60 liver samples with ages ranging from 2 months to 20 months was created based on this same set of 748,955 CpGs. Pearson correlations with age were calculated using the *corrwith* function from the *pandas* package. Linear regressions were computed using the *LinearRegression* function as part of the *sklearn* package.

### Probabilistic single cell clock

To devise an algorithm to ascertain epigenetic age in single cells, we were inspired by recent work from the Wagner lab on analyzing age in individual bisulfite-barcoded-amplicon sequencing (BBA-seq) reads from bulk samples (Han, Franzen, et al., 2020; Han, Nikolić, et al., 2020). First, we calculated linear regressions for every CpG covered in the training dataset in the form:

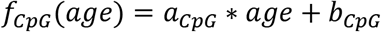

where age in months is the independent variable, *f*_*CpG*_(*age*) is the predicted average methylation level, and *a* and *b* are the linear regression coefficient and intercept, respectively (Fig. 1c). We also calculated the Pearson correlation coefficient with age for every CpG in the training set.

Next, we intersected the CpGs covered in the training dataset with those in any given single cell, producing a series of *n* CpGs that are present in both bulk and individual single-cell profiles (Fig. 1d). We filtered these *n* CpGs based on the absolute value of their correlation with age, selecting (in the liver model) the 700 CpGs with the largest absolute Pearson correlation and (in the multi-tissue model) the 2,000 CpGs with the largest absolute Pearson correlation. These numbers of CpGs to include in each model were determined *in silico* based on those that generated the most optimal accuracy metrics using the Gravina et al. dataset as a benchmark. Of note, diverse numbers of CpGs can be used with minimal fluctuations in epigenetic age predictions (Fig. S5).

For each selected CpG per cell, we iterated through age in steps of 0.1 months from a minimum age to a maximum age parameter. Using the linear regression formula calculated for an individual CpG, we computed *f*_*CpG*_(*age*), which normally lies between 0 or 1. If this value lied outside of the range (0, 1), it was instead replaced by 0.001 or 0.999 depending on the proximity to either value. Next, we assume that the probability of observing a methylated single cell coming from a tissue of a given age is approximately equal to *f*_*CpG*_(*age*), that is, *Pr*_*CpG*_(*age*) = *f*_*CpG*_(*age*). Then, the probability that a single cell is methylated at that CpG is *Pr*_*CpG*_(*age*), and conversely the probability that a single cell is not methylated at that CpG is 1 - *Pr*_*CpG*_(*age*). This provides an age-dependent probability P for every common CpG retained in the algorithm (Fig. 1e)

The product of each of these probabilities will be the overall probability of the observed methylation pattern: 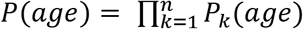 where k represents individual CpGs. Our goal is then to find the maximum of that product among different ages (i.e., to find the most probable age for observing that particular methylation pattern). For that, we took the sum across CpGs of the logarithm of the probabilities (to avoid running into underflow errors during computation). This gives us 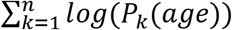 for each age step. These logarithmic sums provide a likelihood metric for every age that a single cell comes from a bulk tissue of that age. Finally, we pick the age of maximum likelihood as our predictor of age for a single cell.

### Computational and statistical analyses

All analyses were conducted using Python 3.8.3 with the standard suite of scientific, mathematical, and plotting packages. Custom bash scripts were used to process sequencing data. Welch’s t-test assuming unequal variances was used to perform all statistical tests. P-values of less than 0.05 were taken as significant.* denotes p < 0.05, ** denotes p < 0.01, and *** denotes p < 0.001.

## ACKNOWLEDGMENTS

We are grateful to Tiamat Fox and Adit Ganguly for help with schematic figures. We also thank Marco Mariotti, Anastasia Shindyapina, Sun Hee Yim, Sang-Goo Lee, Didac Santesmasses and Patrick Griffin for helpful discussion. Supported by NIA grants to VNG.

## AUTHOR CONTRIBUTIONS

AT performed all analyses. AT and CK devised the method and implemented the algorithm. CK contributed to data preprocessing. AT and VNG wrote the manuscript with input from CK. VNG conceived the study and supervised the work.

## CODE AVAILABILITY

The source code for scAge will be made available upon publication.

---

**Figure S1:**
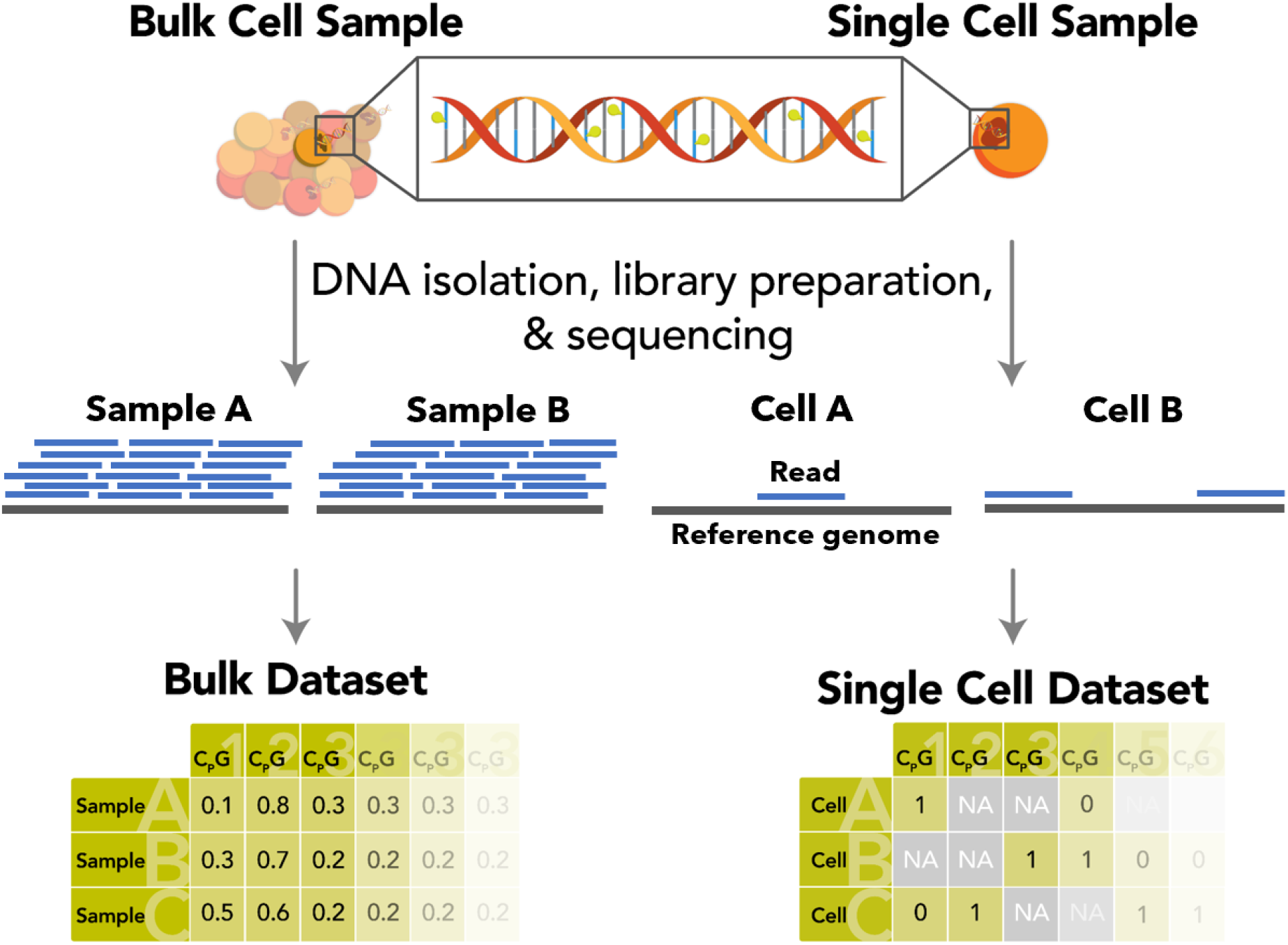
Practical differences in the output of bulk and single cell epigenetic sequencing methods. Schematic comparison of bulk (left) and single cell (right) methylation sequencing approaches. Using bulk samples ensures high and consistent CpG coverage, while single cell profiles typically suffer from effectively random and sparse coverage. Bulk samples produce continuous values from 0 to 1 (due to the presence of reads from different cells), while single cell samples exhibit primarily digitized methylation. Consequently, conventional feature tables (bottom) used to train elastic net regression clocks are unfeasible to create in single cells owing to the presence of extensive missing data.

**Figure S2:**
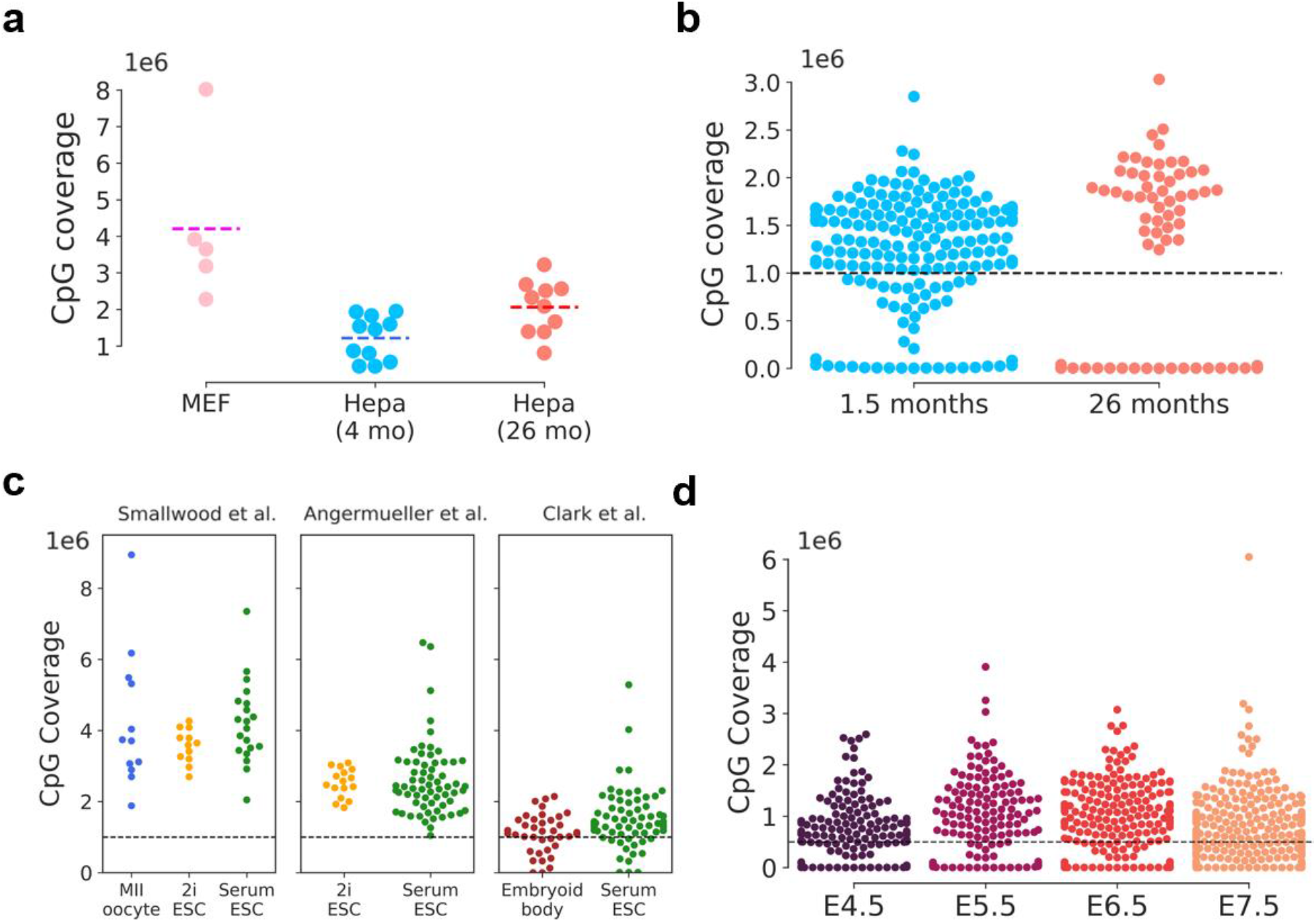
Single-cell CpG coverage varies among datasets. CpG coverage in **(a)** MEFs, 4-month-old and 26-month-old hepatocytes; **(b)** 1.5-month-old and 26-month-old muscle stem cells; **(c)** MII oocytes, ESCs cultured in 2i or serum conditions, and embryoid bodies; and **(d)** E4.5 to E7.5 embryonic cells. Based on the coverage profile and number of cells in each dataset, filtering was applied to remove cells with very low coverage (black dotted lines). No threshold was set in **(a)** due to low cell numbers, a threshold of 1,000,000 CpGs was designated in **(b)** and **(c)**, and a threshold of 500,000 CpGs was set in **(d)** to optimize both prediction consistency and the largest number of cells per study.

**Figure S3:**
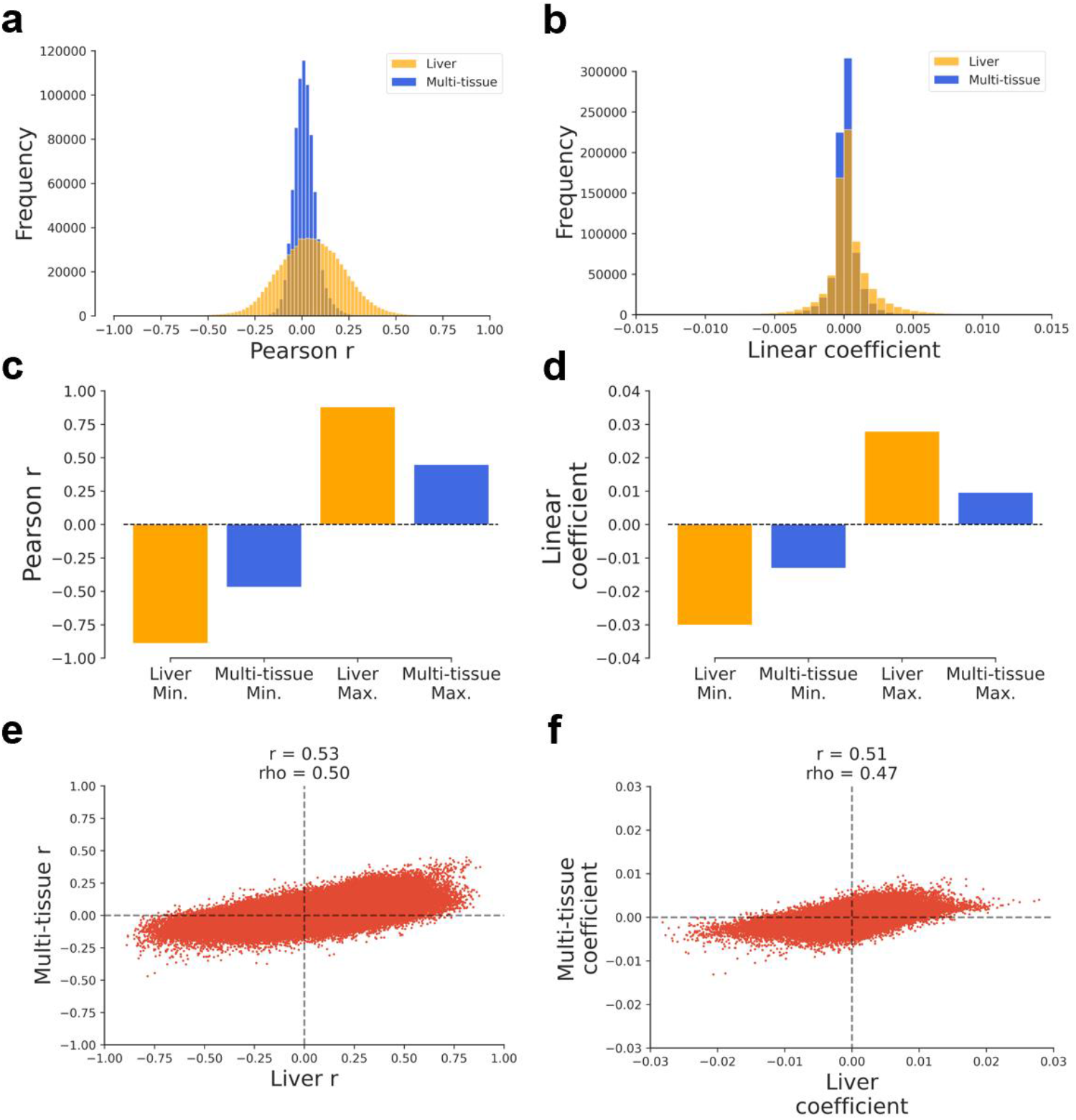
Correlation and regression metrics differ between training datasets. Distribution of Pearson *r* **(a)** and the linear regression coefficients **(b)** for liver and multi-tissue training datasets. Minimum and maximum Pearson *r* **(c)** and linear regression coefficients **(d)** for liver and multi-tissue training datasets. Relationship between multi-tissue and liver training dataset Pearson *r* **(e)** and linear regression coefficients **(f)**.

**Figure S4:**
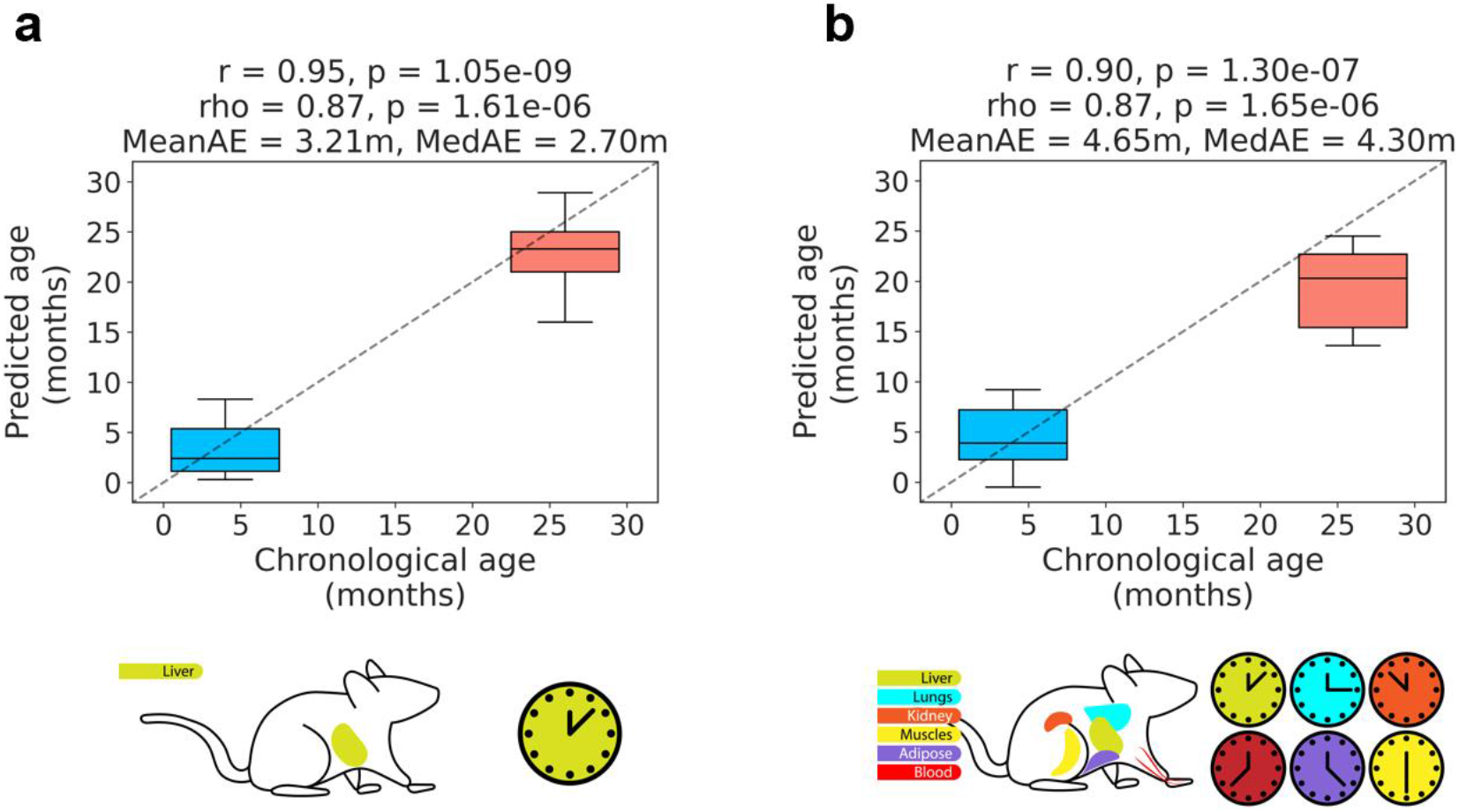
scAge prediction accuracy improves with outlier removal. Epigenetic age predictions on liver hepatocytes with one outlier (accelerated aging) per group removed (SRR3136664 in young and SRR3136629 in old) for both the liver scAge **(a)** and multi-tissue scAge **(b)** models.

**Figure S5:**
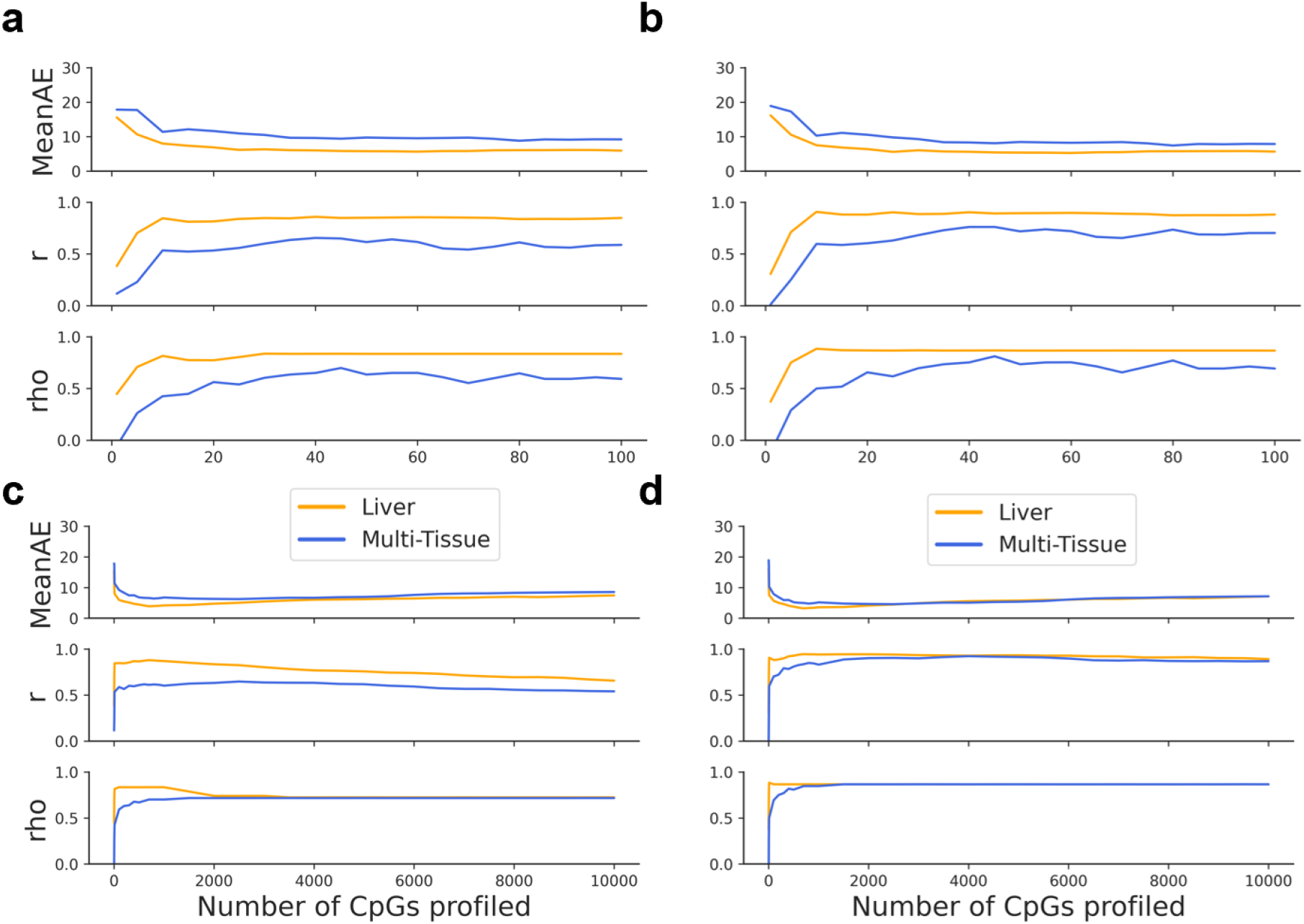
Number of CpGs profiled in scAge affects predictive metrics. scAge predictive metrics (MeanAE: mean absolute error, r: Pearson correlation coefficient, rho: Spearman correlation coefficient) on hepatocytes vary based on the number of CpGs included in the algorithm. Increasing CpGs profiled from 1 to 100 in steps of 5 generally improves prediction accuracy with outliers included **(a)** or with outliers removed **(b)**. Increasing CpGs profiled up to 10,000 per cell in steps of 500 shows that an intermediate number of CpGs results in the highest predictive accuracy, both with outliers included **(c)** and with outliers removed **(d)**.

**Figure S6:**
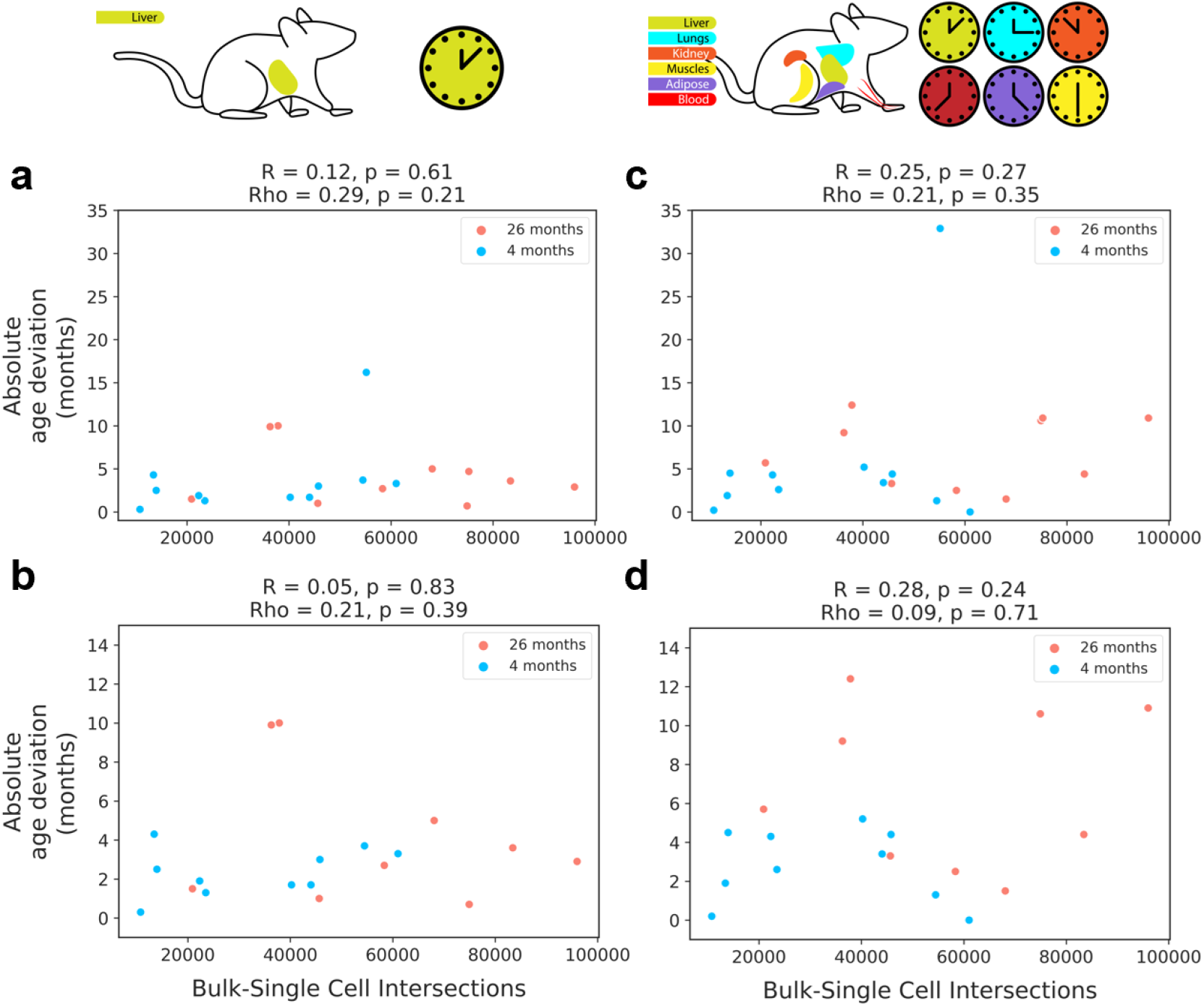
CpG coverage has minimal effect on prediction accuracy. Relationship between the absolute age deviation (absolute value of the difference between predicted age in a single cell and chronological age of the tissue of origin) and the number of common single-cell/bulk CpGs using the liver scAge clock with outliers **(a)** or with the two outliers removed **(b)**, and using the multi-tissue scAge clock with outliers **(c)** or with outliers removed **(d)**.

